# Diagnostic Assessment of Osteosarcoma Chemoresistance Based on Virtual Clinical Trials

**DOI:** 10.1101/022467

**Authors:** K.A. Rejniak, M.C. Lloyd, D.R. Reed, M.M. Bui

## Abstract

Osteosarcoma is the most common primary bone tumor in pediatric and young adult patients. Successful treatment of osteosarcomas requires a combination of surgical resection and systemic chemotherapy, both neoadjuvant (prior to surgery) and adjuvant (after surgery). The degree of necrosis following neoadjuvant chemotherapy correlates with the subsequent probability of disease-free survival. Tumors with less than 10% of viable cells after treatment represent patients with a more favorable prognosis. However, being able to predict early, such as at the time of the pre-treatment tumor biopsy, how the patient will respond to the standard chemotherapy would provide an opportunity for more personalized patient care. Patients with unfavorable predictions could be studied in a protocol, rather than a standard setting, towards improving therapeutic success. The onset of necrotic cells in osteosarcomas treated with chemotherapeutic agents is a measure of tumor sensitivity to the drugs. We hypothesize that the remaining viable cells, i.e., cells that have not responded to the treatment, are chemoresistant, and that the pathological characteristics of these chemoresistant tumor cells within the osteosarcoma pre-treatment biopsy can predict tumor response to the standard-of-care chemotherapeutic treatment. This hypothesis can be tested by comparing patient histopathology samples before, as well as after treatment to identify both morphological and immunochemical cellular features that are characteristic of chemoresistant cells, i.e., cells that survived treatment. Consequently, using computational simulations of dynamic changes in tumor pathology under the simulated standard of care chemotherapeutic treatment, one can couple the pre- and post-treatment morphological and spatial patterns of chemoresistant cells, and correlate them with patient clinical diagnoses. This procedure, that we named ‘Virtual Clinical Trials’, can serve as a potential predictive biomarker providing a novel value added decision support tool for oncologists.

## Introduction

Osteosarcoma (also called *osteogenic sarcoma*) is an aggressive malignant neoplasm arising from osteoblast progenitor. It is the most common primary high grade bone sarcoma. It occurs most often in children and young adults in areas where bone is growing quickly, such as long bones. Osteosarcoma is not a common cancer, with about 800 new cases diagnosed each year in USA; half are in children and teens (1). The diagnosis of this tumor can be usually made on clinical and radiological ground with histological confirmation using the biopsy specimen. Osteosarcoma exhibits a malignancy that produces osteoid matrix. Among various types of osteosarcoma, conventional osteosarcoma is the most common primary osteosarcoma. It is composed of osteoblastic (26-80%), chondroblastic (10-13%) and fibroblastic (10%) variants (2). The standard therapy consists of multi-agent neoadjuvant chemotherapy with doxorubicin and cisplatin often with high dose of methotrexate, followed by surgery and adjuvant chemotherapy with the same agents. With this treatment, the 5-year survival rate for people with localized osteosarcoma is 60-80%. However, if the osteosarcoma has already metastasized, the 5-year survival rate is about 15-30% (1). Immediately after recovery from chemotherapy, patients have their tumors resected and the effect of the chemotherapy on the cancer cells is ascertained. A careful histologic analysis by an experienced pathologist can identify viable osteosarcoma cells, necrotic cells, and other changes, such as fibrosis and hyalinization. According to the Huvos grading system (3), the percentage of necrosis within the tumor tissue determines the tumor response grade and predicts the probability of progression-free survival. Tumors with less than 10% viable cells after chemotherapy (grade III or grade IV response) represent a subset of patients with a more favorable prognosis on the order of 80% 5 year event free survival (EFS) whereas those with greater than 10% viable tumor cells have a similar EFS of closer to 50% (4,5).

Recent efforts in osteosarcoma research have focused on a multinational trial randomizing patients to additional therapy based on histologic necrosis (6). Good risk patients, with less than 10% viable cells, were randomized to receive maintenance pegylated interferon alpha-2b for 74 weeks following completion of standard 6 cycles of doxorubicin, cisplatin, and high dose methotrexate (MAP). Poor risk patients were randomized to additional duration of adjuvant therapy, from 18 weeks to 29 weeks and additional ifosfamide and etoposide in addition to full doses of MAP (7). While not yet published, the presentations have not demonstrated that either investigational arm was superior to standard MAP ((8) and personal communication). Preclinical testing of agents through murine models has identified some agents with promise and has led to the development of one active trial (NCT02097238) in relapsed osteosarcoma patients and as background to several trials in development (9,10). Banking efforts have led to the establishment of a valuable resource for ongoing basic science work (11). Sequencing efforts have illustrated the key role of p53 mutation in this malignancy and investigated the structural variations that characterize this disease (12). Metastatic osteosarcoma continues to be a significant challenge and the most recent clinical trial demonstrated feasibility of adding zoledronic acid to the chemotherapeutic backbone in this setting (13,14). Efforts to better incorporate the young adult population into clinical trials and discern outcomes in this group compared to the younger patients are underway as well (15-17).

There are various histological types of osteosarcoma which have variable clinical behavior from low grade to high grade. Conventional osteosarcoma is the most common type and considered high grade which warrant neoadjuvant chemotherapy. It is known that chondroblastic variant has less therapy effect than osteoblastic and fibroblastic variants (18). However, the current histologic system does not allow for further predictions of patient response and survival probabilities at the time of diagnosis for conventional osteosarcoma in general. Being able to predict early, such as at the time of the tumor pre-treatment biopsy, how the patient will respond to the standard chemotherapy would provide an opportunity for more personalized medical care. For example, this approach would facilitate a trial design with modified therapy for patients with unfavorable features, whereas patients with a predicted positive response could continue to receive the standard-of-care (SOC) treatment or perhaps even less intense therapies.

## Hypothesis

The percentage of viable and necrotic cells in osteosarcomas treated with chemotherapeutic agents is a measure of tumor sensitivity to the drugs. Viable cells that survived the treatment are treatment-resistant (subsequently called *chemoresistant cells*). We hypothesize that virtual assessment of chemoresistance, based on a combination of advanced high content image analyses and feature classification methods applied to patient histology samples, as well as quantitative computational simulations of dynamic changes in tumor pathology during treatment (Virtual Clinical Trials), can predict the responses to the SOC chemotherapeutic treatment of osteosarcoma patients.

## Evaluation of the hypothesis

### Digital Pathology Evaluation

Pathomics is a modern concept and practice that uses computer assistance to perform analysis of pathology images. While high-resolution scanned digital images of patient histology samples are being increasingly used by pathologists in ascertaining tumor grades and assisting in cancer diagnoses, prognoses, and therapy choices, the process of evaluating pathology samples is still traditionally done manually by pathologists equipped with the microscope. However, the computerized analysis of pathology images (pathomics) is slowly but steadily gaining its place in cancer translational research providing a powerful tool to explore the complexities of large and heterogeneous collections of the cells forming the tumor tissue (19,20). Over the last couple of years, several such quantitative methods have been developed for specific clinical applications. For example, a quantitative segmentation scheme EMaGACOR (expectation-maximization driven geodesic active contour, (21,22)) has been designed to detect the extent of lymphocytic infiltration of the breast tumor tissue (that correlates with HER2-positive breast cancer recurrence) from standard H&E staining of histopathology images. These highly effective procedures allow for both the quantification of infiltrating lymphocytes and for the exploitation of the differences in lymphocyte spatial arrangements. A suite of quantitative algorithms has been developed to detect regions containing cancer cells by using the multiresolution digitized images of prostate biopsies or prostatectomies (23-25). In this approach, several images of the same tissue but of different resolutions are used (similarly to the manual approach employed by pathologists), and the most salient features at each resolution are followed and quantitatively classified using appropriate biostatistics methods (boosted Bayesian multiresolution classifier [BBMC] (23), spatially-invariant vector quantization [SIVQ] (24), or probabilistic pairwise Markov models (25)). The machine learning methods have been applied in the Computational Pathologist system (C-Path, (26)) to automatically extract the quantitative morphologic features of both tumor cells and the surrounding stroma from the H&E-stained breast cancer tissue microarrays. As a result, C-Path has identified that features of the stromal tissue adjacent to the cancer were better predictors of patient survival than the features of tumor cells alone. Similar quantitative algorithms have been developed for the automated quantification of IHC-determined protein expressions in melanocytes (27), for the automated assessment of the extent of malignant nuclei in colon cancer histology images (28), and for quantifying the architectural complexity in breast and prostate cancer specimens (29,30). Our group also used quantitative segmentation algorithms to automatically determine scores for HER2-positive, ER-positive, and PR-positive cells within digitized histology images of breast cancer tissues (31), to correlate patient ER statuses with spatial distributions of ER-positive and ER-negative cells, as well as tumor vascular density and vascular area (32), and to determine the extent of the drug-mediated cytotoxic and apoptotic effects in osteosarcoma xenotransplants (33).

There are obvious advantages of having an automated system for digital pathology quantification. Such systems allow for the analysis of a vast amount of data collected from each histology sample. They enable the extraction of accurate, reproducible, quantifiable features of both tumor cells and the stroma. They facilitate computer-aided diagnoses and sharing with multiple people at different locations for remote consultations, tumor boards, or education. There are, however, two drawbacks in current use of digital pathology in clinical practice; one is of a technical nature, the other arises from the data availability. For technical reasons, scanning of histology slides adds an additional time delay in the tissue preparation process. The procedures for image acquisition lack the standardization necessary for automatic quantification. A special IT infrastructure is required to enable the timely accessibility of digital pathology images and their storage. Moreover, the sizes of digitized histology images present a formidable challenge for computer image analyses, both in the number of pixels to analyze (hundreds of millions) and in the time necessary for completing the analysis. All such technical limitations are being addressed by the scanner manufacturers, by the computer system designers, and by bioinformaticians, and it is only a matter of time until faster scanning and analysis systems are available on the market. On the other hand, however, each histological sample, whether assessed and scored manually by a pathologist or automatically by a computerized system, represents data characteristic only for a particular point in the tumor progression (a time of biopsy or tumor recession). These so-called static data provide information neither about how this particular tumor reached the observed state, nor about how this tumor would progress in response to the administered treatment. Obviously, such information could help in determining patient prognosis and in choosing appropriate treatment options. We propose to address this limitation by employing computational simulations (Virtual Pathology) of how a given tumor can respond to a given treatment.

### Computational Simulation Evaluation

Tumor dynamic responses to anti-cancer treatments, both in terms of changes in tumor structure over time and its spatial evolution, can be simulated using quantitative mathematical modeling. When appropriately calibrated, the in-silico models provide a tool to test various scenarios of tumor progression, leading to experimentally testable hypotheses. Some examples of predictive mathematical models include the following. Haeno et al. (34) proposed a mathematical model of metastasis formation calibrated to data from a large series of patients with pancreatic cancers, and predicted both the timing and type of clinical interventions that can most effectively impact patient survival. In particular, the model showed that earlier-applied neoadjuvant chemotherapy provides a significant survival benefit, and neoadjuvant radiation therapy prevents further metastases. Powathil et al. (35) developed a model of cell-cycle phase-specific radiosensitivity that took into account heterogeneity in tumor oxygenation and provided a ranking of different therapeutic regimens in terms of overall treatment efficacy for a given patient. Macklin et al. (36) used patient histopathology data to calibrate a mathematical model of a ductal carcinoma in situ and provided patient-specific quantitative mapping between the calcification geometry observed in mammograms and the actual tumor shape and size offering a tool to more precisely plan for surgical margins. Our group also used mathematical modeling to investigate how the tumor tissue architecture and the extent of extracellular space between the tumor cells influence the transport and distribution of drugs or diagnostic imaging agent molecules inside the tumor (37). This study indicated that for moderately diffusive therapeutic agents, interstitial transport is highly influenced by tumor cell size and packing density, and thus, the morphological features of a given tumor should be considered in determining the therapeutic treatment. In another computational/experimental study we investigated the effects of exogenous pyruvate on transient changes in tumor oxygenation and enhanced effectiveness of hypoxia-activated pro-drugs (HAPs) in hypoxic regions of pancreatic tumors (38). These results suggested that acute increase in tumor hypoxia can improve clinical efficacy of HAPs in pancreatic adenocarcinomas and other tumors. Moreover, we developed a mathematical model incorporating G1/S and G2/M cell-cycle checkpoints (39) to test the effectiveness of cyclin-dependent kinase (CDK) inhibitors on tumor growth-arrest. This in-silico study suggested that when tight tumor cell clusters are exposed to the CDK inhibitors, their growth suppression could be an effect of contact inhibition rather than of the drug mechanism, which may be important in designing the administration schedules and order of inhibitors acting at the G1/S and G2/M cell-cycle checkpoints.

It is evident from the growing literature on mathematical oncology and systems oncology (40-51) that in-silico models are becoming the quantitative tools of choice for understanding the nonlinear interactions between multiple components of complex systems, such as cancer. In particular, these models can address both intracellular and extracellular heterogeneities, as well as dynamic changes in the tissue microenvironment and in tumor responses to therapies. The computational models are capable of quantitatively integrating data from various levels of organization (for example, signaling pathways, cell metabolism, individual or collective cell behavior, tumor microenvironment composition, etc.) into a comprehensive system that can be used to test various therapeutic treatments in a systematic way, as well as to formulate testable treatment hypotheses. Computational models are also flexible and can be refined when additional clinical information becomes available. These tools can easily be used to examine individualized therapies, as, by their nature, the computational models are designed to employ different parameter values and different initial conditions in each execution of the model. The parameters can be derived from patient data, such as biopsies, SOC medical imaging, gene expressions, or proteomics, and are thus patient-specific. However, the main concern that needs to be addressed before such models can be used in clinic is their predictability. This is usually achieved via an iterative cyclic process in which the model is built and validated against the clinical data for which the patients’ outcome is known, and it is refined and crossvalidated again and again until the satisfactory level of accuracy is achieved (this is called ‘a learning phase’). After that the model can be used to make predictions (‘a translational phase’) based on new clinical data and algorithms developed and validated in the learning phase. The important aspect of a clinically relevant computational model is the use of the SOC data that is widely accepted and routinely collected in all clinics. For this reason, we propose to utilize patient biopsy samples, even if they account only for a small subset of a large tumor and may not accurately or consistently represent the tumor as a whole. However, in current clinical practice both tumor diagnosis and therapy choice are based on such a patient’s biopsy sample. Since both a biopsy sample before treatment and a tumor resected after the treatment represents only single time points in tumor development, we use computer simulations to provide a dynamics link between these two static points. By using computer simulations multiple times, we can determine the most possible paths in tumor progression and calculate the likelihood of osteosarcoma chemoresistance. Such computer simulations of temporal and spatial changes in tumor composition after exposure to various therapeutic options can provide a clinician with a decision support tool, the Virtual Clinical Trials.

## Plan for hypothesis validation

We propose here the Virtual Clinical Trials procedure for determining the chemoresistance of a given osteosarcoma based on the data extracted from a patient’s biopsy. The learning phase of our predictor algorithm (Fig. 1 A-F) will use retrospective data from patients of a known progression-free survival status. Patient histopathology samples before and after treatment will be compared using advanced image analyses (pathomics), feature classification methods (biostatistics and morphometry), and computational simulations of tumor progressions (virtual pathology). This will lead to the identification of patterns of cellular features that are characteristic of chemoresistant osteosarcoma cells (the cells that survived the therapy). Once validated, this predictor will be used for prospective studies (the translational phase of the predictor, Fig. 1 I-IV). In this case, the pre-treatment biopsy samples will be analyzed and compared with previously identified features of chemoresistant cells, as well as with the previously created library of the simulated results that determine tumor response to treatment. This will allow to calculate the likelihood of the given tumor being chemoresistant. The details of the construction of our predictor (both phases) are given below.

**Fig 1.**
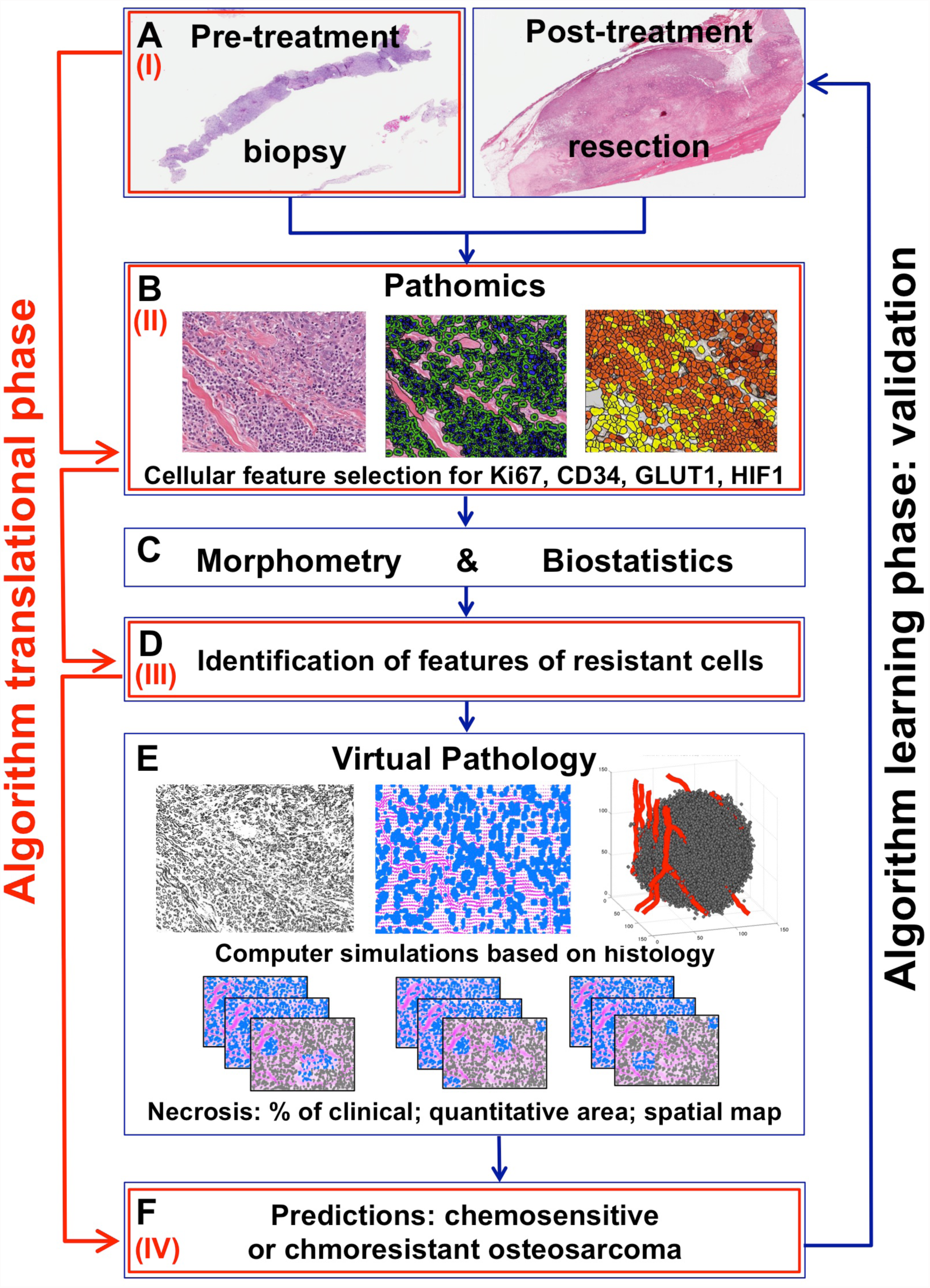
Schematic of a Virtual Clinical Trial Predictor of Tissue Chemoresistance. *Learning phase:* **A** Pre- and post-treatment data; **B** Pathomics: feature selection and analysis; **C** Biostatistics and morphometry analysis; **D** Identification of features of chemoresistant cells; **E** Virtual Pathology: simulations and predictive reports; **F** Likelihood of tumor tissue chemoresistance; *Translational phase:* **(I)** Collection of pre-treatment data; **(II)** Pathomics analysis; **(III)** Comparison with previously identified features of chemoresistant cells; **(IV)** Comparison with virtual pathology reports for predicting the likelihood of a tumor tissue being chemoresistant.

### Learning phase Fig. 1A, Pre- and post-treatment tissues

Pre-treatment biopsy samples and post-treatment tissue resection samples from patients that responded well to SOC chemotherapy (<10% of cells are viable), and from poorly responding patients (>10% of cells are viable) will be selected. All tissue slices will be stained with H&E, as well as with IHC markers for proliferation (Ki67), hypoxia-inducible factor (HIF-1), glucose transporter (GLUT-1), tumor protein p53, and endothelial cells (CD34). The 40× magnification with 0.25μm/pixel resolution slide scans will be taken for advanced image analysis.

### Learning phase Fig. 1B, Analysis of digital images of tumor tissue

High-resolution digitized images of tumor histology will be analyzed using advanced image analysis techniques (Pathomics) to identify and quantify morphological and immunochemical features of individual tumor cells. First, the regions of interest from the whole tumor tissue image will be determined, and the spatial zones of a low-to-high intensity of staining are selected; then, the segmentation of individual cell nuclei and cell cytoplasms will be performed, and both the physical and molecular features in each individual cell will be extracted. The physical features of each cell will include morphological or textural parameters, such as and nuclei size, shape, compactness, density, and the cytoplasm to nucleus ratio. The molecular futures evaluated from the IHC-stained slices will include cell and cytoplasm staining intensity for each individual cell, as well as the localization of tumor tissue vasculature. These will be used in the quantitative classification.

### Learning phase Fig. 1C, Quantitative analysis of cellular features

Quantitative multi-parametric feature sets extracted from individual tumor cells will be collected and used to detect and characterize cells that cluster around particular phenotypes (chemoresistant or chemosensitive). Of particular interest is whether cells will be uniformly distributed in the multi-dimensional morphospace, or whether they will form discrete and detectable sub-populations that cluster around particular morphological archetypes (morphotypes). This analysis will allow for the consideration of multiple features of millions of individual cells that otherwise are difficult to assess visually.

### Learning phase Fig. 1D, Identification of features of chemoresistant cells

Comparison of the quantitative features of individual osteosarcoma cells before and after the treatment will lead to the identification of features that are characteristic of the cells that remained viable after chemotherapy, which are considered *chemoresistant*. Similarly, a set of cellular features that are present in tumors prior to treatment but not after treatment will be considered typical of *chemosensitive* cells.

### Learning phase Fig. 1E, Computer simulations

To determine the likelihood of osteosarcoma tumors being chemoresistant based solely on their pre-treatment histology, multiple computer simulations will be run using patient’s histology data (Virtual Pathology) to provide dynamic response of this specific osteosarcoma tumor to SOC chemotherapeutic treatment. The model will be calibrated to reflect patient-specific features of chemosensitive and chemoresistant cells, as described above. The vasculature identified by CD34 will act as a source of chemotherapeutic agents, oxygenation and nutrients; Ki67 staining determines the tumor proliferative index and defines the initial population of proliferating cells; HIF1a and GLUT1 will definite the cellular metabolic state. Different runs of the model will account for heterogeneous cell-cell and cell-microenvironment interactions, and the percentage of viable cells after each simulation will be counted and reported together with tumor morphology (chemo-charts).

### Learning phase Fig. 1F, Predictions

An analysis of simulation outcomes, such as the extent of tumor necrosis in terms of the necrotic ratio relative to the tumor area, the quantitative area of detectable necrosis and a spatial necrosis map, and the spatial localization of remaining viable (chemoresistant) cells, will determine the likelihood of a patient tumor being chemoresistant (calculated as a percentage of computer simulations classified as chemoresistant). For the learning stage, model feasibility will be tested by comparing model simulations with patient post-treatment histologies, and by comparing with its clinical classifications into chemoresistant or chemosensitive (learning phase, Fig. 1 A–F).

### Translational phase Fig. 1 I–IV

In the translational phase, pre-treatment data (Fig. 1-I) will be used to perform a pathomics analysis (Fig. 1-II), as well as to compare to the features of chemoresistant cells already identified during the learning stage (Fig. 1-III). Then, information on the tumor tissue morphological structure and information on the identified set of cellular features will be used to map these parameters on the chemo-charts created during the learning stage. This will allow for the determination of the likelihood of tumor tissue chemoresistance (Fig. 1-IV). The simulated results can support both sarcoma pathologists and sarcoma clinicians in grading osteosarcomas, making therapeutic decisions and in clinical trial design.

## Consequences of the hypothesis and discussion

We propose here that the Virtual Clinical Trials, a combination of a quantitative analysis of patient histology samples and a predictive modeling of tumor tissue response to treatments, can be used to predict whether a given patient’s tumor will be sensitive to the SOC chemotherapeutic treatment. Having such a high content analysis system that can be assessed at diagnosis will aid in a better prognostic ability. For example, patients with unfavorable predictions from the Virtual Clinical Trials system could be randomized to receive a modified therapy critical for therapeutic success. We are aware that tumor heterogeneity exists in osteosarcoma. While, in our practice, the pre-treatment biopsy specimens from osteoblastic and fibroblastic variants of osteosarcoma were representative of the post-treatment resection specimens, further studies are needed to examine this heterogeneity in more detail. We based our Virtual Clinical Trial concept on the SOC clinical data and clinical practice where therapeutic decisions are based upon examining patients’ biopsy data. It is worth noticing, that recently patients were randomized based on the onset of necrosis (the current best predictive biomarker) for a clinical trial conducted through a multinational collaboration from 2003-2011. Unfortunately the experimental arms of this study have been presented in a preliminary form without demonstrating an improvement over doxorubicin, cisplatin, and high dose methotrexate. However, a computer-assisted analysis of tumor cellular features, such as the one described in this paper, has the potential to be translated to the clinic to aid in the prediction of osteosarcoma response to SOC treatment. This system will improve osteosarcoma trial development, and it can be used to provide an accurate objective means for assessing the morphological characteristics of osteosarcomas, leading to the objective stratification of patients. Ultimately, the Virtual Clinical Trials system can have a translational potential for improving patient care and may be incorporated into the pathologist and clinician’s decision-supporting toolboxes.

